## Supplementary information for "Naïve individuals promote collective exploration in homing pigeons"

### Methods

**Relative position of birds over time.** We determined the relative position of each bird within the pair using the distance $d_{EF}$ of the experienced bird from the naïve one projected onto the direction of motion of the flock. We modified the method proposed in (Nagy et al. 2010), which gives the distance of the experienced bird from the centre of the flock, to also include the segment from the centre of the flock to the naïve bird. This is given by

$$d_{EN}(i) = \left(\overrightarrow{x_E}(i) - \overrightarrow{x_N}(i)\right) \cdot \overrightarrow{v_{pair}}(i) \cdot 2,$$

where $\overrightarrow{x_E}(i)$ and $\overrightarrow{x_N}(i)$ are the positions of the experienced and the naïve bird and $\overrightarrow{v_{pair}}(i)$ is the normalized velocity of the pair. The normalized velocity is computed as

$$\overrightarrow{v_{pair}}(i) = \frac{\langle \overrightarrow{\dot{x}_k}(i) \rangle_k}{\left| \langle \overrightarrow{\dot{x}_k}(i) \rangle_k \right|}.$$

As the flock is composed of two birds only, the projected distance of the experienced bird from the naïve one projected onto the direction of motion of the pair is positive, when the experienced bird is flying *ahead* of the naïve one, and it is negative when the experienced bird is flying *behind*.

**Exploration–exploitation with respect to the previous generation.** Our primary approach to define exploration and exploitation is based on comparing pairs of successive releases using the trajectory of the previous release as our baseline model of exploitation and then labelling newer portions of a focal route as exploration. The model of exploitation (i.e., baseline trajectory) thus varies for each focal release similarly to a moving-average window over successive releases. An alternative approach to define exploration and exploitation is to consider a constant model of exploitation for each release within a given generation. This can be obtained by setting the baseline trajectory to equal the last trajectory of the previous generation. As a consequence, exploitation is defined on the basis of the information available only to the experienced bird at the beginning of a new generation; every portion of the route within that generation that is more than 300 meters away from the baseline is considered exploration.

At the first generation of the experimental group, when birds are trained individually for the successive transmission chain experiment and there is no previous generation available to provide us with a baseline trajectory, we consider the first release as the baseline trajectory for the remainder of the generation. In the case of solo and fixed pairs controls, for which there is no obvious definition of a generation, we considered the 60 releases in each of these experiments as formed by 5 generations, each lasting 12 releases, and used an equivalent approach to that of the experimental group to define exploration and exploitation.

### Results

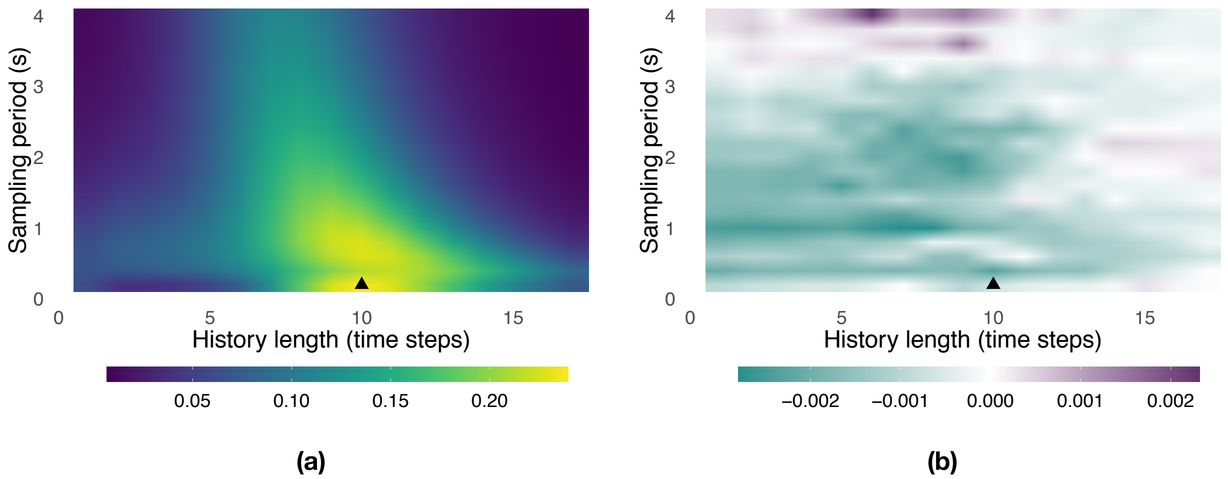


*Figure S1. Landscape of information transfer as a function of the history length, $k \in \{1, \ldots, 17\}$, and of the sampling period, $\{0.2, 0.4, \ldots, 4.0\}$ seconds. Panel (a) shows the total transfer of information between the pair of birds, $T_{E\to N} + T_{N\to E}$, averaged over all releases and generations. Panel (b) shows the net transfer of information between the pair of birds, $T_{E\to N} - T_{N\to E}$, averaged over all releases and generations. Positive (respectively, negative) values represent configurations where the experienced (naïve) bird is more informative than the naïve (experienced) one. The triangle represents the configuration with maximum total transfer of information.*

**Landscape of information transfer and choice of parameters.** We first explored how information transfer between the birds varied as a function of two parameters: the sampling period (seconds) and the history length used in the computation of transfer entropy. Our original GPS data are sampled at a frequency of 1 sample every 0.2 seconds (5 Hz); we can further subsample these data by dropping samples, for example, using one sample every 0.4, 0.6, ..., 4.0 seconds as the sampling period. The history length represents the number of past rotations by the bird we want to predict that we consider when computing transfer entropy from the other bird in the pair.

The total transfer of information between the birds varies as a function of these parameters (Figure S1a). It peaks in the region delimited by history lengths of 8–10 time-steps and a sampling period between 0.2–1.2 seconds, whereas it vanishes otherwise. The total transfer of information, which is generally adopted as a measure to choose study parameters (Porfiri 2018), reaches its maximum for a history length of $k = 10$ at one sample every 0.2 seconds (black triangle in Figure S1a). We use this parameter configuration for the rest of our information-theoretic analysis. For this and for most other parameter configurations, the naïve bird is more informative about the future behaviour of the experienced one than the other way around (Figure S1b). Only in two regions, both far from the point maximizing the total transfer of information, is the experienced bird more informative than the naïve one; however, the overall amount of information transferred between birds in these regions is negligible.

**Comparison of information transfer with the surrogate dataset.** To verify their significance, we compared our estimates of information transfer between the two birds with equivalent measurements taken from the surrogate dataset. We expect causal effects measured in the original dataset to be stronger than those found in the surrogate one. As shown in Table S1, our expectations are fully met: the original dataset shows values of transfer entropy significantly higher than those observed in the surrogate dataset, both for the entire dataset as well as for each separate generation.

*Table S1. Statistical comparison of information transfer between the original and the surrogate dataset over all generations and over separate generations. Column 1 reports the generation and sample sizes. Columns 2 and 4 report the differences between the mean value of transfer entropy of the original dataset and that of the surrogate dataset. Columns 3 and 5 report the results of one-sided two-sample Whitney–Mann–Wilcoxon rank-sum tests with continuity correction ($p$-value and $W$ statistic) testing if the original dataset has significantly higher transfer entropy than the surrogate one. Significant $p$-values are reported in bold.*

| | *Original vs surrogate dataset* | | | |
|---|---|---|---|---|
| *Generation* | $T_{E\to N} - T^s_{E\to N}$ | $H_1: T_{E\to N} > T^s_{E\to N}$ | $T_{N\to E} - T^s_{N\to E}$ | $H_1: T_{N\to E} > T^s_{N\to E}$ |
| All ($n = 343, n^s = 29035$) | $\mu = 0.0089$ | $\boldsymbol{p < .001}$ ($W = 6145522$) | $\mu = 0.0088$ | $\boldsymbol{p < .001}$ ($W = 6126284$) |
| 2 ($n = 94, n^s = 7912$) | $\mu = 0.0062$ | $\boldsymbol{p < .001}$ ($W = 445262$) | $\mu = 0.0074$ | $\boldsymbol{p < .001}$ ($W = 452733$) |
| 3 ($n = 99, n^s = 9801$) | $\mu = 0.0094$ | $\boldsymbol{p < .001}$ ($W = 615721.5$) | $\mu = 0.0119$ | $\boldsymbol{p < .001}$ ($W = 645665.5$) |
| 4 ($n = 81, n^s = 6561$) | $\mu = 0.0071$ | $\boldsymbol{p = .006}$ ($W = 308433.5$) | $\mu = 0.0057$ | $\boldsymbol{p = .015}$ ($W = 303085.5$) |
| 5 ($n = 69, n^s = 4761$) | $\mu = 0.0111$ | $\boldsymbol{p < .001}$ ($W = 214258.5$) | $\mu = 0.0075$ | $\boldsymbol{p = .002}$ ($W = 197618.5$) |

**Analysis of time spent by each bird at the front of the pair.** We divided each flight into different segments, with each segment representing a consecutive portion of the route with either the experienced or the naïve bird at the front of the pair, and then measured the segment durations (Figure S2). The distribution of segment durations resembles a log-normal distribution for both the entire dataset and for individual generations. Experienced and naïve birds are characterized by very similar distributions closely overlapping each other. Overall, the naïve bird spent significantly longer periods of time at the front of the pair (Table S2), but the difference is small and largely driven by that of generation 3 (the only generation showing significant differences). With the exception of generation 3, the experienced and the naïve bird spent, on average, an approximately equal portion of the route at the front of the pair (Table S3). At generation 3, the naïve bird was at the front of the pair for a significantly larger portion of the route with respect to the experienced bird (respectively, 54% versus 46%).

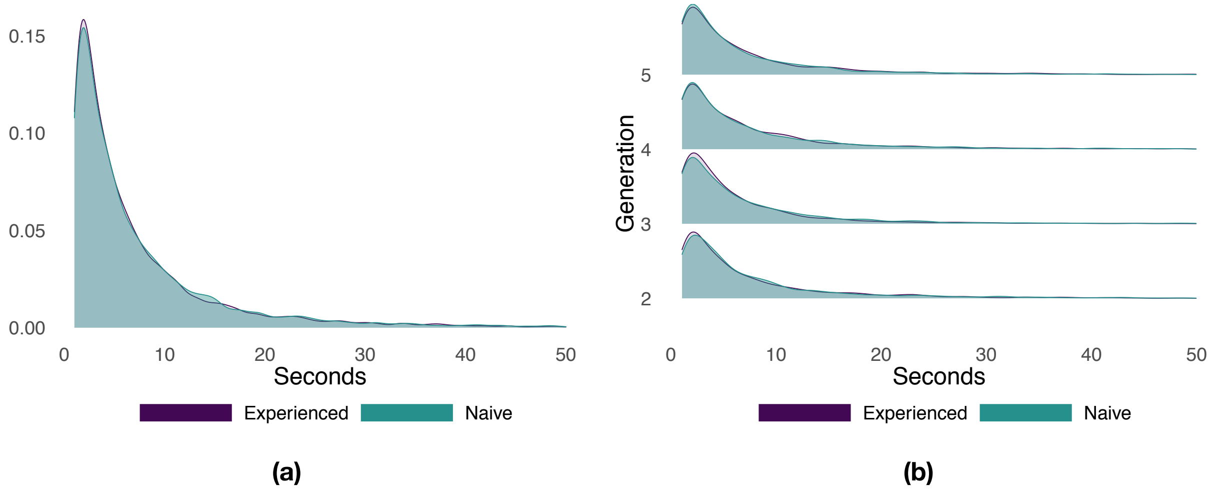


*Figure S2. Probability density function of the duration of flight segments with either the experienced or the naïve bird at the front of the pair. Panel (a) shows the results aggregated over all generations. Panel (b) shows the results separately for each generation.*

*Table S2. Statistics for the duration of flight segments with either the experienced or the naïve bird at the front of the pair. Column 1 reports the generation and sample sizes. Columns 2 and 3 give the mean duration and the standard deviation of segments for the experienced and for the naïve bird. Column 4 reports the results of two-sided two-sample Whitney–Mann–Wilcoxon rank-sum tests with continuity correction ($p$-value and $W$ statistic) testing differences between the distribution of the duration $D$ of flight segments for the two birds. Significant $p$-values are reported in bold.*

| *Generation* | *Experienced* | *Naïve* | $H_1: D_E \neq D_N$ |
|---|---|---|---|
| All ($n^E = 10725, n^N = 10773$) | $\mu = 7.93, \sigma = 12.92$ | $\mu = 8.23, \sigma = 14.12$ | $\boldsymbol{p = .047}$ ($W = 56865638$) |
| 2 ($n^E = 2512, n^N = 2492$) | $\mu = 8.89, \sigma = 17.51$ | $\mu = 8.81, \sigma = 13.55$ | $p = .13$ ($W = 3052262$) |
| 3 ($n^E = 3690, n^N = 3737$) | $\mu = 6.83, \sigma = 9.08$ | $\mu = 7.96, \sigma = 14.51$ | $\boldsymbol{p < .001}$ ($W = 6588674$) |
| 4 ($n^E = 2297\ n^N = 2283$) | $\mu = 8.63, \sigma = 13.65$ | $\mu = 8.55, \sigma = 14.23$ | $p = .86$ ($W = 2630132$) |
| 5 ($n^E = 2226, n^N = 2261$) | $\mu = 7.98, \sigma = 11.23$ | $\mu = 7.73, \sigma = 13.93$ | $p = .23$ ($W = 2568854$) |

*Table S3. Statistics for the proportion of a flight with either the experienced or the naïve bird at the front of the pair. Column 1 reports the generation and sample size. Columns 2 and 3 give the mean and the standard deviation of the proportion of a flight with either the experienced or the naïve bird at the front of the pair. Column 4 reports the results of two-sided paired Wilcoxon signed-rank tests with continuity correction ($p$-value and $V$ statistic) testing differences between the distribution of the proportion $P$ of a flight for the two birds. Significant $p$-values are reported in bold.*

| *Generation* | *Experienced* | *Naïve* | $H_1: P_E \neq P_N$ |
|---|---|---|---|
| All ($n = 341$) | $\mu = 0.49, \sigma = 0.2$ | $\mu = 0.51, \sigma = 0.2$ | $p = .46$ ($V = 27817.5$) |
| 2 ($n = 92$) | $\mu = 0.51, \sigma = 0.23$ | $\mu = 0.49, \sigma = 0.23$ | $p = .69$ ($V = 2243$) |
| 3 ($n = 99$) | $\mu = 0.46, \sigma = 0.17$ | $\mu = 0.54, \sigma = 0.17$ | $\boldsymbol{p = .03}$ ($V = 1851$) |
| 4 ($n = 81$) | $\mu = 0.5, \sigma = 0.2$ | $\mu = 0.5, \sigma = 0.2$ | $p = .84$ ($V = 1705$) |
| 5 ($n = 69$) | $\mu = 0.49, \sigma = 0.2$ | $\mu = 0.51, \sigma = 0.2$ | $p = .95$ ($V = 1219$) |

**Exploration versus exploitation with respect to successive releases.** We used a distance-based mechanism to label portions of a focal route as either exploration or exploitation depending on the point-to-point distance from each point of the focal route to the closest point of the baseline route at the previous release. To determine a suitable threshold, we compared successive trajectories flown by the experienced bird towards the end of training (generation 1, last three flights) and studied the distribution of distances between successive trajectories (Figure S3a). The distribution of distances is right-skewed and approximately exponential. A threshold of 300 meters captures a large portion of the probability mass, about 70.5%, whereas larger distances are progressively less likely. On this basis, we set a threshold of 300 meters to distinguish between phases of exploitation ($< 300$ meters) and phases of exploration ($\geq 300$ meters). Figure S3b shows the distribution of distances between consecutive flights observed during each generation whereas Figure S4a shows the same results aggregated over all generations with details of the proportion of time  that each bird is leading as a function of the same distance showed in Figure S4d. Figures S4b and S4c show similar results for fixed pairs of birds and solo individuals. Whereas fixed pairs are characterized by a distribution similar to that of experimental pairs, solo birds have a markedly shifted distribution towards exploitation at the expense of exploration.

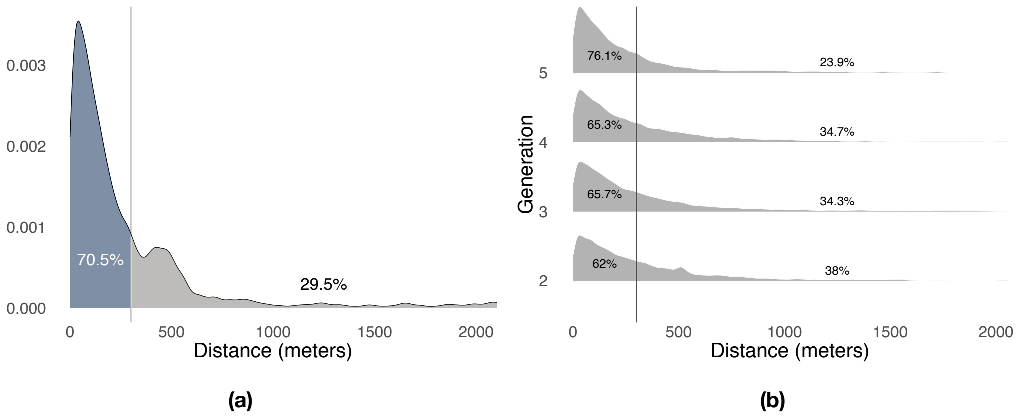


*Figure S3. Distribution of minimum distances between the pairs of consecutive flights. Panel (a) shows the results for the trained birds during the first generation of the experiment. Panel (b) shows the results for the pair of birds during all remaining generations of the experiments. Vertical lines highlight the division of the probability mass between exploitation (left) and exploration (right) defined by a 300 meters threshold.*

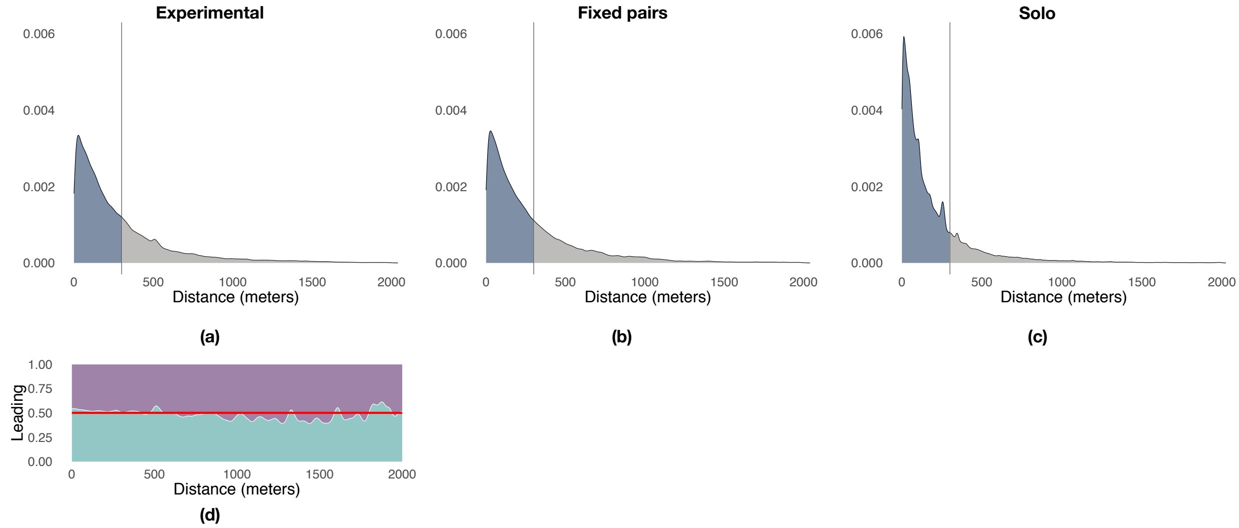


*Figure S4. Illustration of the distribution of point-to-point distances between pairs of consecutive flights highlighting the 300 meter threshold that demarks the end of exploitation and the beginning on exploration. Panel (a) reports the results for experimental pairs over generations 2–5, panel (b) reports those for the fixed pairs control, and panel (c) those for the solo control. Panel (d) shows the proportion of times experienced (purple) and naïve (green) birds are leading the flock as a function of the distance between the current and the previous trajectory. The red line indicates an equal likelihood between the two birds to lead the flock.*

We compared the proportion of exploration and exploitation across experimental conditions before the beginning (*i.e.*, generation 1) and during transmission chain experiments (*i.e.*, generation 2 to 5). During the first 12 releases (Table S4), individuals from the experimental group that flew solo during generation 1 could not be distinguished from birds in the solo control whereas fixed pairs of birds showed levels of exploration significantly higher than those of solo birds. During releases 13 to 60, that is, when naïve individuals are iteratively introduced in the transmission chains at the beginning of each generation, experimental pairs showed instead significantly higher levels of exploration than both solo and fixed pairs of birds, with these former exploring much less than birds that flew in pairs.

*Table S4. Statistical comparison of mean proportions of a flight spent exploring versus exploiting across treatments (experimental pairs, solo and fixed pairs controls) for the first 12 releases (generation 1) and for releases 13 to 60 (generation 2 to 5). Entries report the proportion of exploration vs exploitation for pairs of treatments as well as the results of two-sided two-sample Whitney–Mann–Wilcoxon rank-sum tests with continuity correction ($p$-value and $W$ statistic) for differences in proportion of exploration. Significant $p$-values are reported in bold. Results of testing for the proportion of exploitation are equivalent and not repeated below.*

| *Releases* | *Dataset* | Solo control | Fixed pairs control |
|---|---|---|---|
| 1 to 12 | Experimental (gen. 1) | Row: 36.7% vs 63.3%<br>Col: 34.2% vs 65.8%<br>$p = .55$ ($W = 5288$) | Row: 36.7% vs 63.3%<br>Col: 51.7% vs 48.3%<br>$\boldsymbol{p < .001}$ ($W = 2230$) |
| | Solo control | – | Row: 36.7% vs 63.3%<br>Col: 34.2% vs 65.8%<br>$\boldsymbol{p < .001}$ ($W = 1837$) |
| 13 to 60 | Experimental (gen. 2-5) | Row: 32.9% vs 67.1%<br>Col: 15.7% vs 84.3%<br>$\boldsymbol{p < .001}$ ($W = 94108.5$) | Row: 32.9% vs 67.1%<br>Col: 29.3% vs 70.7%<br>$\boldsymbol{p = .0456}$ ($W = 50472$) |

| | | | |
|---|---|---|---|
| | Solo control | – | Row: 15.7% vs 84.3%<br>Col: 29.3% vs 70.7%<br>$\boldsymbol{p < .001}$ $(W = 31517)$ |

**Exploration versus exploitation with respect to the previous generation.** In addition to the exploration–exploitation analysis with respect to successive releases, we also performed a similar analysis where exploitation is defined with respect to the previous generation (see supplementary Methods above) in order to validate the robustness of the main results obtained with our primary approach. The distributions of point-to-point distances between baseline and focal trajectories (Figure S5) resembled those observed when comparing successive releases (Figure S4). Differences across experimental and control treatments (Table S5) were much more pronounced under this model but in line with the results obtained above (Table S4). Moreover, the overall exploration trends reported in Figure S6 showed the same signatures of the analysis over successive releases (Figure 5a). Exploration decreased over generations in all experimental conditions with experimental pairs exploring more than both fixed pairs of birds and solo individuals.

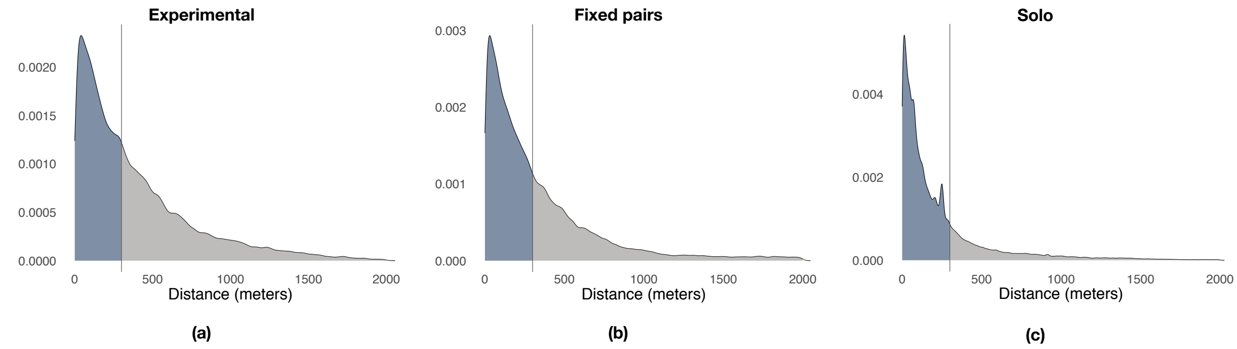


*Figure S5. Illustration of the distribution of point-to-point distances between each flight of a generation (focal trajectories) and the last flight of the previous generation (baseline trajectory). Colors and vertical lines highlight the 300 meter threshold that demarks the end of exploitation and the beginning of exploration. Panel (a) reports the results for experimental pairs over generations 2–5, panel (b) reports those for the fixed pairs control, and panel (c) those for the solo control.*

*Table S5. Statistical comparison of mean proportions of a flight spent exploring versus exploiting across treatments (experimental pairs, solo and fixed pairs controls) for the first 12 releases (generation 1) and for releases 13 to 60 (generation 2 to 5) when considering the last release at the previous generation as the baseline trajectory. Entries report the proportion of exploration vs exploitation for pairs of treatments as well as the results of two-sided two-sample Whitney–Mann–Wilcoxon rank-sum tests with continuity correction ($p$-value and $W$ statistic) for differences in proportion of exploration. Significant $p$-values are reported in bold. Results of testing for the proportion of exploitation are equivalent and not repeated below.*

| *Releases* | *Dataset* | Solo control | Fixed pairs control |
|---|---|---|---|
| 1 to 12 | Experimental (gen. 1) | Row: 61.5% vs 38.5%<br>Col: 55.8% vs 44.2%<br>$p = .11$ $(W = 5592)$ | Row: 61.5% vs 38.5%<br>Col: 82.9% vs 17.1%<br>$\boldsymbol{p < .001}$ $(W = 1721)$ |

| | | | |
|---|---|---|---|
| | Solo control | – | Row: 55.8% vs 44.2%<br>Col: 82.9% vs 17.1%<br>$\boldsymbol{p < .001}$ ($W = 1163$) |
| 13 to 60 | Experimental (gen. 2-5) | Row: 46.6% vs 53.4%<br>Col: 18.9% vs 81.1%<br>$\boldsymbol{p < .001}$ ($W = 103095$) | Row: 46.6% vs 53.4%<br>Col: 32.4% vs 67.6%<br>$\boldsymbol{p < .001}$ ($W = 60502$) |
| | Solo control | – | Row: 18.9% vs 81.1%<br>Col: 32.4% vs 67.6%<br>$\boldsymbol{p < .001}$ ($W = 30744$) |

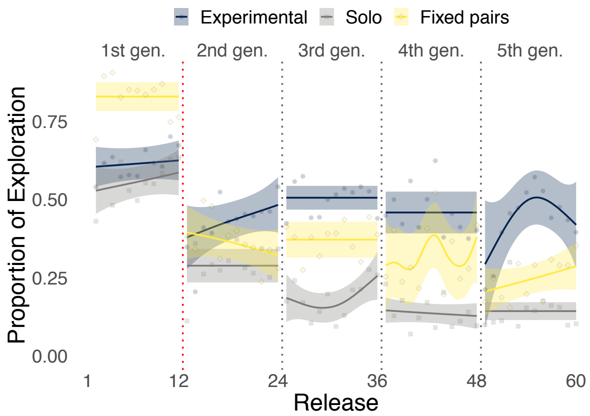


*Figure S6. Illustration of the proportion of exploration (respectively, one minus the proportion of exploitation) over releases when considering the last release at the previous generation as the baseline trajectory. Results are shown for the experimental group, the solo control, and the fixed pairs control. The red dotted vertical line delineates the end of the experimental group's training phase. Smoothed lines are computed with generalized additive models using shrinkage cubic regression splines (mean and standard error); points represent averages for individual releases.*

**The role of leadership during exploration and exploitation.** Using this distance-based mechanism, we label each segment of a flight as either exploration or exploitation. For each segment, we keep track of which bird, experienced or naïve, is leading the pair. We then compute the proportion of time spent by the experienced bird and by the naïve one leading the pair during either an exploration or exploitation phase (Table S6). The combined proportion of time of both birds in either exploration or exploitation gives instead the corresponding contribution of the pair. Overall, each bird spends approximately 16.5% of the time leading the pair during an exploration phase and 33.5% of the time leading the pair during an exploitation phase. With the exception of exploitation phases in generation 3, there is no significant difference between the proportion of time spent by the two bird in an exploration or exploitation phase (Table S7).

*Table S6. Proportion of a flight led by each of the two birds, calculated separately for exploration and exploitation phases. Column 1 reports the generation and sample size. Columns 2 and 3 give the mean and the standard deviation of the proportion of a flight led, respectively, by the experienced and the naïve bird for the case of exploration. Columns 4 and 5 give the mean and the standard deviation of the proportion of a flight led, respectively, by the experienced and the naïve bird for the case of exploitation.*

| | *Exploration* | | *Exploitation* | |
|---|---|---|---|---|
| *Generation* | *Experienced* | *Naïve* | *Experienced* | *Naïve* |
| All ($n = 341$) | $\mu = 0.17, \sigma = 0.15$ | $\mu = 0.16, \sigma = 0.13$ | $\mu = 0.32, \sigma = 0.18$ | $\mu = 0.35, \sigma = 0.2$ |

| | | | | |
|---|---|---|---|---|
| 2 $(n = 92)$ | $\mu = 0.2, \sigma = 0.17$ | $\mu = 0.17, \sigma = 0.14$ | $\mu = 0.31, \sigma = 0.2$ | $\mu = 0.32, \sigma = 0.21$ |
| 3 $(n = 99)$ | $\mu = 0.16, \sigma = 0.13$ | $\mu = 0.19, \sigma = 0.13$ | $\mu = 0.3, \sigma = 0.16$ | $\mu = 0.36, \sigma = 0.17$ |
| 4 $(n = 81)$ | $\mu = 0.18, \sigma = 0.16$ | $\mu = 0.16, \sigma = 0.13$ | $\mu = 0.32, \sigma = 0.18$ | $\mu = 0.34, \sigma = 0.22$ |
| 5 $(n = 69)$ | $\mu = 0.13, \sigma = 0.13$ | $\mu = 0.1, \sigma = 0.1$ | $\mu = 0.37, \sigma = 0.17$ | $\mu = 0.4, \sigma = 0.21$ |

*Table S7. Statistical comparison of leadership by experienced vs. naïve birds, tested separately for exploration and exploitation over all generations and over separate generations. Column 1 reports the generation and sample size. Columns 2 and 3 report the results of two-sided paired Wilcoxon signed-rank tests with continuity correction ($p$-value and $V$ statistic) for differences in proportion of flight led between experienced and naïve birds for exploration and exploitation, respectively. Significant $p$-values are reported in bold.*

| | *Experienced vs Naïve* | |
|---|---|---|
| *Generation* | *Exploration* | *Exploitation* |
| All $(n = 341)$ | $p = .44\ (V = 28820)$ | $p = .13\ (V = 26240)$ |
| 2 $(n = 92)$ | $p = .17\ (V = 2490)$ | $p = .71\ (V = 2048)$ |
| 3 $(n = 99)$ | $p = .08\ (V = 1891)$ | $\boldsymbol{p = .035}\ (V = 1871)$ |
| 4 $(n = 81)$ | $p = .36\ (V = 1767)$ | $p = .79\ (V = 1603)$ |
| 5 $(n = 69)$ | $p = .22\ (V = 1187)$ | $p = .8\ (V = 1131)$ |

**Transitions between phases of exploration and exploitation.** Although once initiated, phases of exploration and phases of exploitation are led in equal manner by the experienced and the naïve bird, the experience unbalance within the pair might affect the likelihood of a bird to initiate transitions from one phase of the exploration–exploitation process to other. To investigate this question, we measured transition probabilities for both birds over generations (Table S8) and over releases (Table S9). Transitions from exploration to exploitation are not significantly different between experienced and naïve birds, but transitions from exploitation to exploration are significantly (albeit marginally) more likely to be initiated by naïve birds. However, this result seems driven by the data of generation 3, where the naïve individual is marginally but significantly more likely than the experienced one to initiate exploration phases, while in all other generations both birds are equally likely to initiate changes in either direction. When looking at transition probabilities over releases, we found a similar trend except for the first release where the naïve bird is much more likely to initiate exploration phases.

*Table S8. Statistical comparison of the proportion of transitions from exploitation to exploration and from exploration to exploitation led by the experienced and by the naïve bird over generations. Column 1 reports the generation number. Columns 2 and 4 report the estimated probabilities that transitions are led by the experienced bird, $P_E$, and by the naïve bird, $P_N$, for transitions, respectively, from exploitation to exploration and from exploration to exploitation. Columns 3 and 5 give the results of exact binomial tests ($p$-value and sample size $n$) of the null hypothesis that the probability $P_E$ that transitions are led by the experienced bird equals 0.5. Significant $p$-values are reported in bold.*

| | *Exploitation → Exploration* | | *Exploration → Exploitation* | |
|---|---|---|---|---|
| *Generation* | $P_E$ *versus* $P_N$ | $H_1: P_E \neq 0.5$ | $P_E$ *versus* $P_N$ | $H_1: P_E \neq 0.5$ |
| All | $P_E = 0.467, P_N = 0.533$ | $\boldsymbol{p = .042}\ (n = 964)$ | $P_E = 0.513, P_N = 0.487$ | $p = .42\ (n = 966)$ |
| 2 | $P_E = 0.494, P_N = 0.506$ | $p = .9\ (n = 247)$ | $P_E = 0.52, P_N = 0.48$ | $p = .56\ (n = 244)$ |

| | | | | |
|---|---|---|---|---|
| 3 | $P_E = 0.432, P_N = 0.568$ | $\mathbf{p = .02}$ $(n = 301)$ | $P_E = 0.483, P_N = 0.517$ | $p = .6$ $(n = 300)$ |
| 4 | $P_E = 0.5, \quad P_N = 0.5$ | $p = 1$ $(n = 216)$ | $P_E = 0.539, P_N = 0.461$ | $p = .28$ $(n = 219)$ |
| 5 | $P_E = 0.45, P_N = 0.55$ | $p = .18$ $(n = 200)$ | $P_E = 0.522, P_N = 0.478$ | $p = .57$ $(n = 203)$ |

*Table S9. Statistical comparison of the proportion of transitions from exploitation to exploration and from exploration to exploitation led by the experienced and by the naïve bird over releases. Column 1 reports the release number. Columns 2 and 4 report the estimated probabilities that transitions are led by the experienced bird, $P_E$, and by the naïve bird, $P_N$, for transitions, respectively, from exploitation to exploration and from exploration to exploitation. Columns 3 and 5 give the results of exact binomial tests ($p$-value and sample size $n$) of the null hypothesis that the probability $P_E$ that transitions are led by the experienced bird equals 0.5. Significant $p$-values are reported in bold.*

| | *Exploitation → Exploration* | | *Exploration → Exploitation* | |
|---|---|---|---|---|
| *Release* | $P_E$ *versus* $P_N$ | $H_1: P_E \neq 0.5$ | $P_E$ *versus* $P_N$ | $H_1: P_E \neq 0.5$ |
| 1 | $P_E = 0.295, P_N = 0.705$ | $\mathbf{p < .01}$ $(n = 44)$ | $P_E = 0.477, P_N = 0.523$ | $p = .88$ $(n = 44)$ |
| 2 | $P_E = 0.478, P_N = 0.522$ | $p = .81$ $(n = 69)$ | $P_E = 0.522, P_N = 0.478$ | $p = .81$ $(n = 67)$ |
| 3 | $P_E = 0.519, P_N = 0.481$ | $p = .83$ $(n = 81)$ | $P_E = 0.575, P_N = 0.425$ | $p = .22$ $(n = 80)$ |
| 4 | $P_E = 0.576, P_N = 0.424$ | $p = .27$ $(n = 66)$ | $P_E = 0.6, P_N = 0.4$ | $p = .14$ $(n = 65)$ |
| 5 | $P_E = 0.487, P_N = 0.513$ | $p = .91$ $(n = 76)$ | $P_E = 0.519, P_N = 0.481$ | $p = .82$ $(n = 77)$ |
| 6 | $P_E = 0.54, P_N = 0.46$ | $p = .48$ $(n = 100)$ | $P_E = 0.465, P_N = 0.535$ | $p = .55$ $(n = 99)$ |
| 7 | $P_E = 0.443, P_N = 0.557$ | $p = .34$ $(n = 88)$ | $P_E = 0.494, P_N = 0.506$ | $p = 1.0$ $(n = 89)$ |
| 8 | $P_E = 0.409, P_N = 0.591$ | $p = .18$ $(n = 66)$ | $P_E = 0.597, P_N = 0.403$ | $p = .14$ $(n = 67)$ |
| 9 | $P_E = 0.438, P_N = 0.562$ | $p = .22$ $(n = 112)$ | $P_E = 0.496, P_N = 0.504$ | $p = 1$ $(n = 115)$ |
| 10 | $P_E = 0.422, P_N = 0.578$ | $p = .17$ $(n = 90)$ | $P_E = 0.522, P_N = 0.478$ | $p = .75$ $(n = 92)$ |
| 11 | $P_E = 0.391, P_N = 0.609$ | $p = .053$ $(n = 87)$ | $P_E = 0.453, P_N = 0.547$ | $p = .45$ $(n = 86)$ |
| 12 | $P_E = 0.541, P_N = 0.459$ | $p = .52$ $(n = 85)$ | $P_E = 0.482, P_N = 0.518$ | $p = .83$ $(n = 85)$ |
